# Competitive mapping allows to identify and exclude human DNA contamination in ancient faunal genomic datasets

**DOI:** 10.1101/2020.03.05.974907

**Authors:** Tatiana R. Feuerborn, Elle Palkopoulou, Tom van der Valk, Johanna von Seth, Arielle R. Munters, Patrícia Pečnerová, Marianne Dehasque, Irene Ureña, Erik Ersmark, Vendela Kempe Lagerholm, Maja Krzewinska, Ricardo Rodríguez-Varela, Anders Götherström, Love Dalén, David Díez-del-Molino

## Abstract

**Background:** After over a decade of developments in field collection, laboratory methods and advances in high-throughput sequencing, contamination remains a key issue in ancient DNA research. Currently, human and microbial contaminant DNA still impose challenges on cost-effective sequencing and accurate interpretation of ancient DNA data.

**Results:** Here we investigate whether human contaminating DNA can be found in ancient faunal sequencing datasets. We identify variable levels of human contamination, which persists even after the sequence reads have been mapped to the faunal reference genomes. This contamination has the potential to affect a range of downstream analyses.

**Conclusions:** We propose a fast and simple method, based on competitive mapping, which allows identifying and removing human contamination from ancient faunal DNA datasets with limited losses of true ancient data. This method could represent an important tool for the ancient DNA field.

## Background

Right after the death of an organism, microbial communities colonize the decomposing tissues and together with enzymes from the organism they start degrading the DNA molecules (Lindahl 1993; Renaud, Schubert, et al. 2019). DNA degradation is dependent on time and environmental variables such as temperature but also humidity and acidity (Kistler et al. 2017). Even though the specific model for DNA decay is still debated and it is likely multifactorial (Kistler et al. 2017), the consequence is that ancient remains typically contain very few quantities of endogenous DNA and these sequences are characterized by short fragment sizes (Dabney et al. 2013).

A second major challenge of ancient DNA research is contamination from exogenous sources (Malmström et al. 2005; Clio Der Sarkissian et al. 2015). Environmental DNA molecules in the soil matrix the ancient sample was recovered from can easily overwhelm the small amounts of endogenous DNA of the ancient sample. This is also true for DNA from people who collected and handled the samples in the field and/or museum collections (C. Der Sarkissian et al. 2014; Green et al. 2006). While the use of Polymerase Chain Reaction (PCR) technology allowed ancient DNA research to overcome these low concentration problems, the sensitivity of the PCR has made it very difficult to avoid introducing modern contaminant sequences among the authentic ancient DNA (Willerslev and Cooper 2005).

In the last decade, together with more refined DNA extraction and laboratory methods tailored to efficiently retrieve very short and scarce DNA sequences (Gamba et al. 2016; Dabney et al. 2013), it has become possible to obtain massive amounts of sequences from ancient material using high-throughput sequencing technologies. These technologies have allowed the recovery of hundreds of ancient human (reviewed in Slatkin and Racimo (2016) and other high quality ancient faunal genomes such as those from horses (Orlando et al. 2013), wooly mammoths (Palkopoulou et al. 2015), and bears (Barlow et al. 2020). However, the challenges from exogenous contamination remain and have sparked a search for computational methods to identify and monitor contaminant DNA sequences in ancient sequencing datasets.

Aside from the short fragment size, the other most notable characteristic of ancient DNA is post-mortem damage. After death, the repairing mechanisms of DNA damage such as hydrolysis and oxidation stop functioning, and this damage accumulates in predictable patterns (Renaud, Schubert, et al. 2019). The most common ancient DNA damage is deamination of cytosines to uracils in the overhangs of fragmented DNA molecules (Gilbert et al. 2003; Stiller et al. 2006; Briggs et al. 2007). This results in an excess of C to T substitutions in the 5’ end (and G to A in the 3’ end) of ancient DNA sequences. Since this feature is very common in sequences derived from ancient DNA sources and absent in younger samples, it has been widely used as a key criteria to authenticate ancient DNA experiments (Dabney et al. 2013; Sawyer et al. 2012).

In modern-day ancient DNA studies, exogenous sequences are differentiated from real ancient sequences from the source organism by mapping all sequences to a reference genome and keeping only those that result in alignments with less than a defined number of differences (Prüfer et al. 2010; Kircher 2012). This approach to circumvent environmental contamination has gained general acceptance, and currently exogenous contaminants are at most considered problematic due to their consumption of sequencing capacity. However, the probability of spurious alignments from exogenous sequences occurring by chance increases with decreasing sequence length (Smith, Waterman, and Burks 1985). In order to avoid these, thresholds for minimum fragment length, that still allow for enough specificity of the alignments, are used (Green et al. 2010; Matthias Meyer et al. 2016; de Filippo, Meyer, and Prüfer 2018).

Modern human contamination is especially problematic for human palaeogenomic studies since ancient, anatomically modern humans typically fall within the variation of modern humans (Allentoft et al. 2015; Lazaridis et al. 2014). This has led to the development of a plethora of methods aimed at computationally quantifying and monitoring exogenous contamination in ancient human DNA datasets (Matthias Meyer et al. 2012; Fu et al. 2014; Rasmussen et al. 2015; Racimo, Renaud, and Slatkin 2016). However, the number of methods that allow for the effective exclusion of this type of contamination remains limited. For example, (Skoglund et al. 2014) used the differential empirical distributions of post-mortem damage (PMD) scores, based on both base quality scores and their level of polymorphism with respect to the reference genome, to differentiate DNA sequences from ancient and modern samples. The PMD scores in a contaminated ancient sample could then be used to successfully identify and separate the sequences that are most likely to have originated from an ancient template molecule from the contaminant ones. Even though this method can allow for the enrichment of the proportion of ancient sequences several-fold in respect to the contaminant sequences, the amount of data lost in the process is very large (45%-90% depending on the age of the ancient sample, Skoglund et al. 2014).

Here we investigate the presence of exogenous sequences in ancient sequencing files to evaluate the pervasiveness of human contamination in ancient faunal DNA studies. We use competitive mapping to identify the levels of contamination in ancient faunal sequencing files and characterize the exogenous sequences by using summary statistics to compare them to those of authentic ancient DNA. We then present this strategy as a simple and fast method that enables the conservative removal of human contamination from ancient faunal datasets with a limited loss of true ancient DNA sequences.

## Results

We first mapped the raw reads from all sequenced samples (50 ancient dogs, *Canis lupus familiaris*, and 20 woolly mammoths, *Mammuthus primigenius*) to three separate reference genomes: the African savannah elephant, dog and human. We found variable levels of sequences confidently mapped to foreign reference genomes (average 0.25% for non-target and 0.86% human) in these sequencing files (Fig. 1A). Most of the files (>95%) contained less than 0.071% of sequences mapped to human and 0.054% the non-target species. We then estimated average read length (mRL) and post-mortem damage scores (PMD^R^) for all alignments and detected significant differences in both indices between sequences mapping to target and to non-target and human, but not between the sequences mapping to the non-target species and human references (Fig. S1).

**Figure 1.**
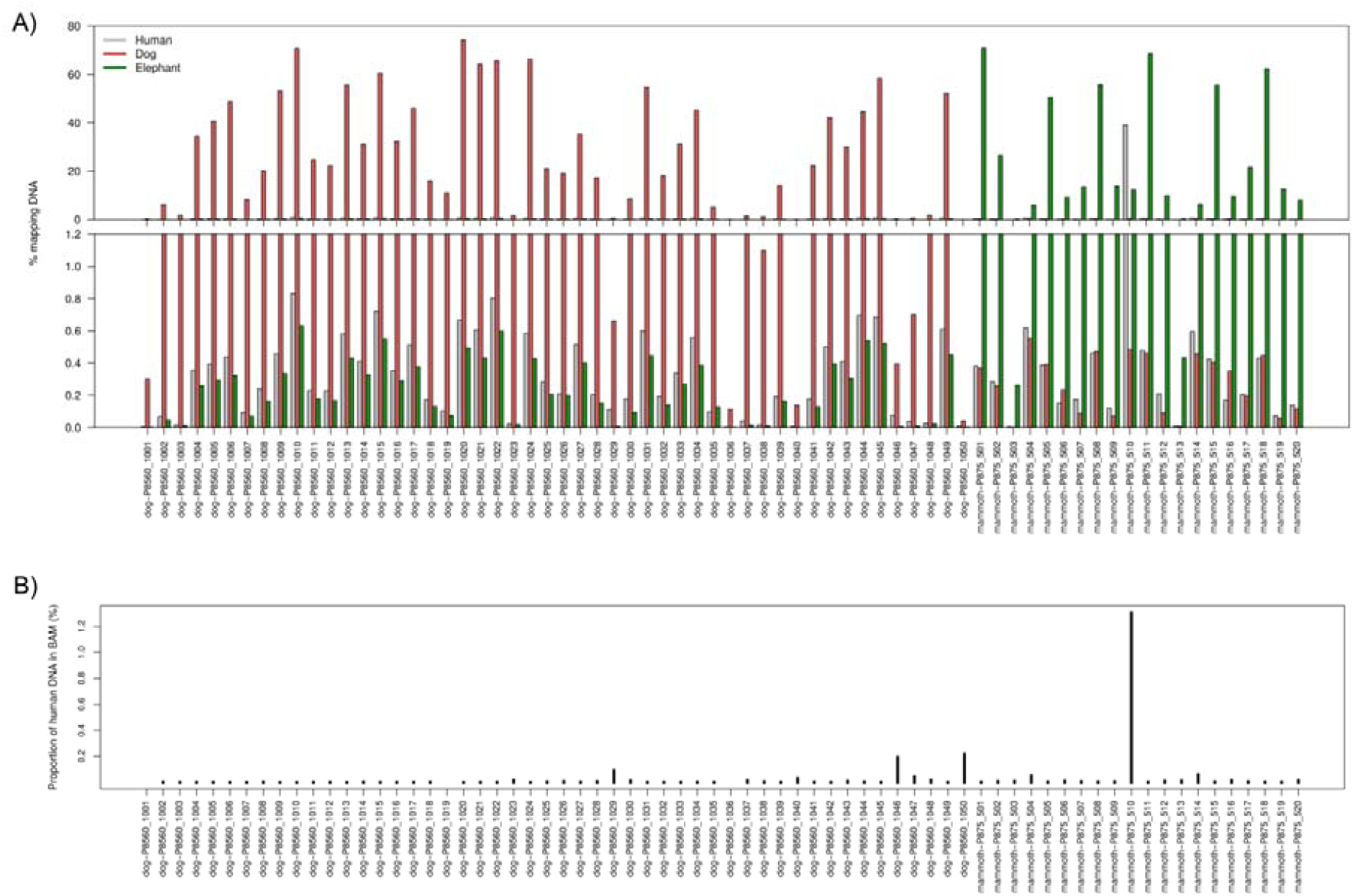
Mapping statistics for target, non-target and human references. A) Upper panel, percentage of reads from each sample mapping to each of the three reference genomes. Lower panel, same as before but zoomed to percentages below 1.2%. B) Proportion of reads from the faunal BAM file that mapped to the human part of the concatenated reference genome.

To investigate whether the target BAM files contain human contaminant sequences we remapped the aligned reads to a concatenated reference composed by the reference genome of the target species, dog or elephant, and the human reference genome (Fig. 2A). This *competitive mapping* approach allowed us to differentiate between three kinds of reads contained in the target species BAM files. First, reads which align to the target reference genome and not to the human reference genome. These sequences represent the endogenous alignments that originate from the sample and not from human or microbial contamination. Second, reads which align to the human reference genome and not to the target species reference genome. These sequences represent the fraction of human contamination in the faunal BAM files. And third, reads that align to both the target reference and the human reference genomes. These sequences could have three origins, 1) true endogenous sequences from regions of the genome highly conserved or identical to the human genome, 2) human contaminant sequences from regions of the genome highly conserved or identical to the target genome, or 3) microbial contaminant sequences that would align to any mammalian genome by random chance. In any case, because these sequences map to both target and human reference genomes at the same time they would thus be discarded when applying mapping quality filters (Fig. 2A).

**Figure 2.**
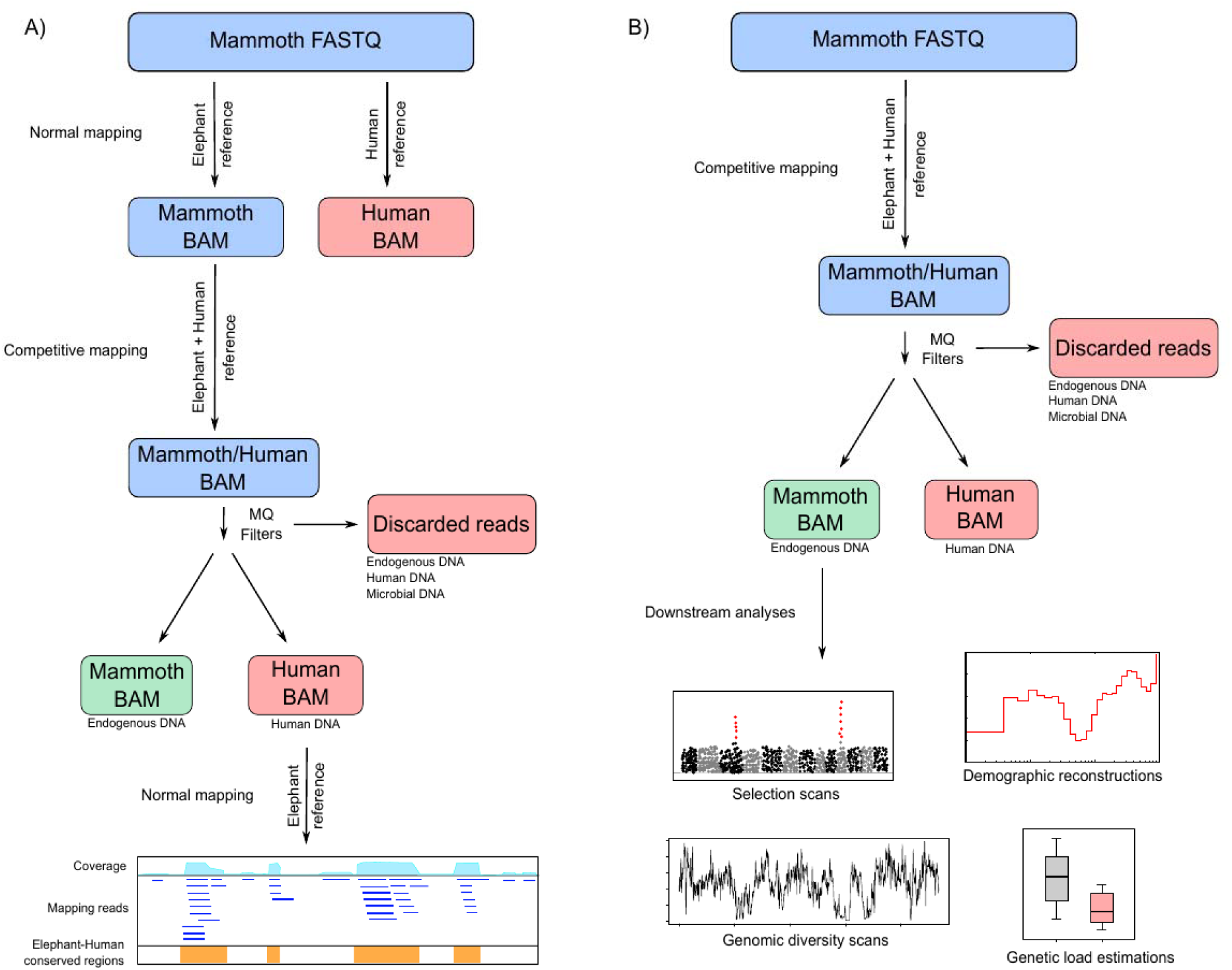
Schematic view of the competitive mapping analyses. FASTQ files represent ‘raw’ sequencing files and BAM files represent alignments to a reference genome. Color boxes indicate different types of data: blue, files that need processing; red, discarded data; and green, data for downstream analyses. A) Schematic view of the analyses performed in this manuscript. An example using a mammoth sample is shown. First, normal mapping to the elephant reference. Second, competitive mapping to a concatenated reference of an elephant and human to detect human contamination in the alignments. Third, normal mapping human data to the elephant reference to check that they map preferentially to conserved regions of the genome. B) Schematic view of a competitive mapping pipeline using a mammoth sample. After competitive mapping, only the sequences mapping to the elephant part of the concatenated reference will be used for downstream analyses.

**Figure 3.**
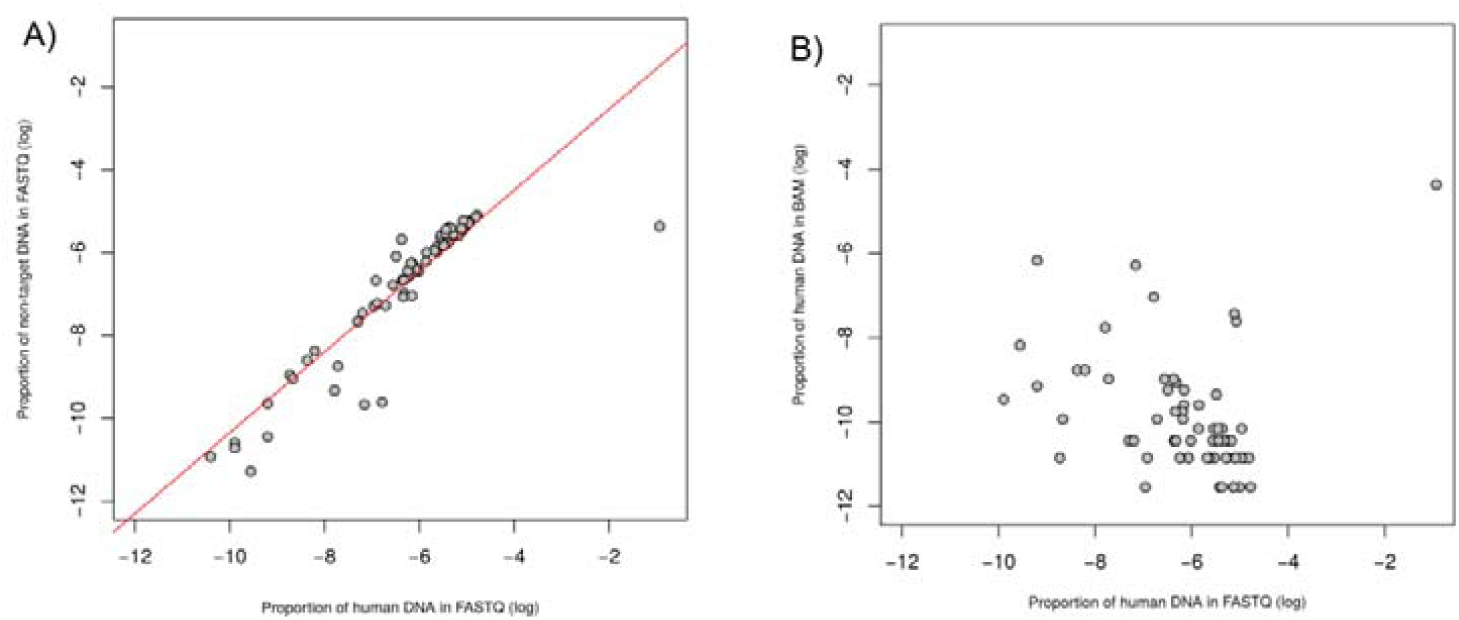
Proportions of sequences mapping to human, target and non-target reference from the FASTQ and BAM files. A) Correlation between the proportion of reads mapping to human and to the non-target species in the raw FASTQ sequencing files (r^2^ = 0.81, F= 303.8, p-value= <2.2e-16). B) Correlation between the proportion of reads mapping to human in the raw FASTQ sequencing files and the proportion of reads mapping to human from the faunal BAM file (r^2^ = 0.01, F= 1.7, p-value= 0.2).

For each sample, we extracted the reads aligned to the target species of the concatenated reference, representing the true ancient sequences, as well as the human, representing the amount of human contamination contained in the original target BAM file. We found that the alignment files from almost all samples contained sequencing reads that preferentially mapped to the human part of the reference genome than to the target part (average 0.03%; range 0 – 1.3%) (Fig. 1B, Supplementary Table 1). However, we caution that, because an unknown fraction of the reads discarded due to the mapping quality filters should also be human contaminant, the fraction of reads in the human part of the concatenated reference represents only a lower bound for the amount of contamination in the original faunal BAM file. Finally, both mRL and PMD^R^ were significantly lower in the sequences mapped to the human part than in the ones mapped to the target (Fig. 4).

**Figure 4.**
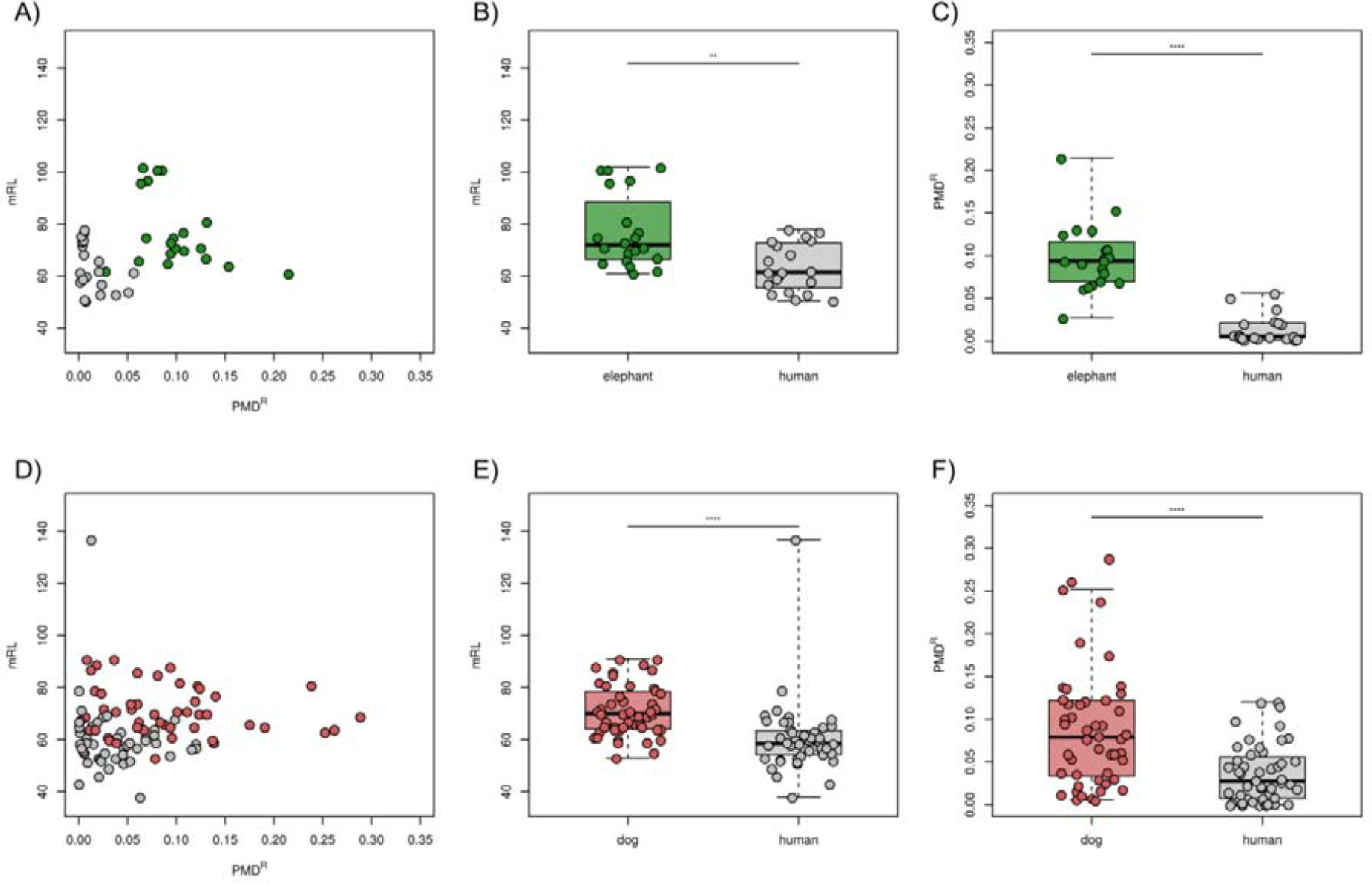
Characterization of endogenous and human contaminant reads in faunal BAM files. A) Comparisons of PMD^R^ and mRL for all mammoth samples. B) mRL for mammoth sequences mapping to the elephant or the human parts of the concatenated reference (Wilcoxon rank sum test, W= 313.5, p-value = 0.00223). C) PMD^R^ for mammoth sequences mapping to the elephant or the human parts of the concatenated reference (Wilcoxon rank sum test, W = 397, p-value = 1.016e-10). D) Comparisons of PMD^R^ and mRL for all ancient dog samples. E) mRL for dog sequences mapping to the dog or the human parts of the concatenated reference (Wilcoxon rank sum test, W = 1929, p-value = 1.251e-08). F) PMD^R^ for dog sequences mapping to the dog or the human parts of the concatenated reference (Wilcoxon rank sum test, W = 1743, p-value = 1.511e-05). In all cases, **: p-value < 0.01 and ****: p-value < 0.0001.

When using competitive mapping, a fraction of sequences that align to both the target and the human parts of the concatenated reference, were lost (Fig. 2A). Our results indicated that this fraction was an average of 1.33% of the total number of reads per sample (range 0.6 – 4.3%, Fig. 5, Supplementary Table 1). However, when accounting only for conserved regions between the target species genome and the human genome, the amount of lost sequences was higher (average 4.53%; range 2.7 – 17.8%).

**Figure 5.**
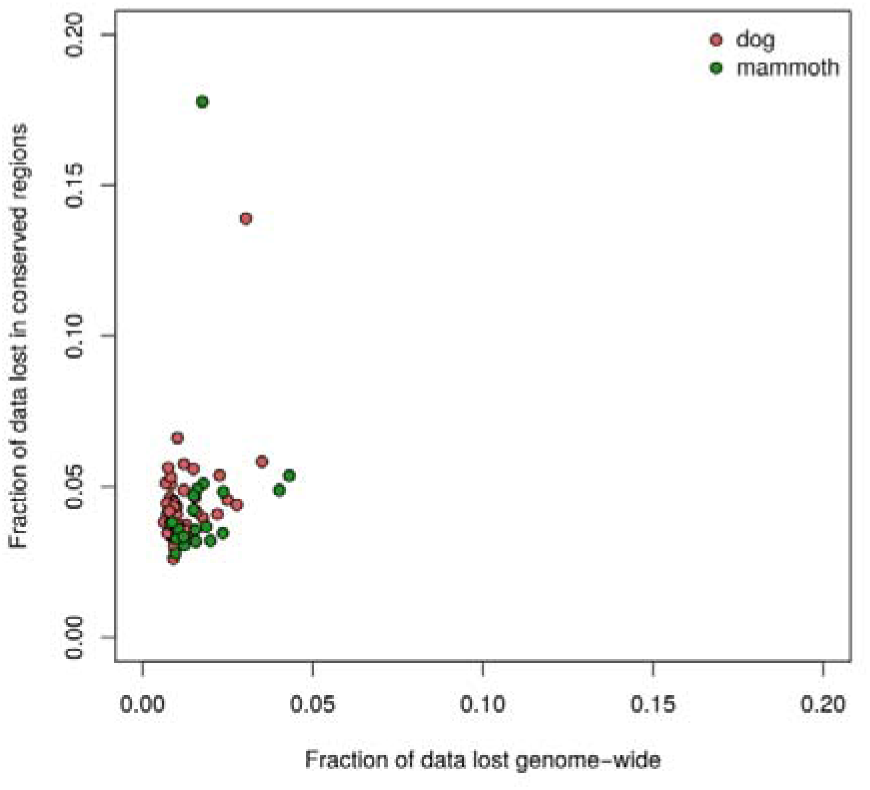
Data lost per sample after competitive mapping. Fraction of data lost in each sample at genome-wide level and only in conserved regions. Colors indicate different species.

## Discussion

### Contamination in raw sequencing files

Overall, we found low levels of sequences mapped to foreign reference genomes in the raw sequencing files (Fig. 1A). In fact, the proportion of reads mapping to the non-target species and human were highly correlated (Fig. 3A), suggesting that they had a common origin. Given that human DNA is a common mammal contamination source in ancient DNA studies (Malmström et al. 2005; Hofreiter, Serre, et al. 2001; Cooper and Poinar 2000; Korlević et al. 2015), it is then likely that a variable amount of contaminant human reads map to the two reference genomes used here, elephant and dog. In fact, there were notable exceptions to the amount of faunal sequences mapping to human, for example one sample contained a higher proportion of sequences mapped to the human (38.9%) than to the target species (12.3%). This suggested that there could be high levels of human DNA contamination in particular sequencing files.

When characterizing mRL and PMD^R^ in the sequences mapping to the different reference genomes we found differences between the sequences mapping to target compared to non-target and human (Fig. S1), in line with the latter being mostly composed by contaminant sequences and the former mostly true endogenous reads. Interestingly, our results suggest almost no differences between the sequences mapping to the non-target species and human references, reinforcing the idea that these two files are composed of sequences with a common origin.

### Human contamination in faunal BAM files

Given that we detected contaminant human sequences in all our ancient fauna sequencing files, we next used competitive mapping to explore whether these contaminant reads can be also found in the BAM file of the target species that would be used for downstream genomic analyses. We found that the BAM files from almost all samples contained sequencing reads that preferentially mapped to the human part of the concatenated reference genome, but the proportion was generally low (Fig. 1B). Interestingly, the proportion of reads mapped to the human reference from the raw data and the fraction of reads mapping to the human part of the concatenated reference in the target BAM after competitive mapping are not correlated (Fig. 3B). This indicates that the amount of human contamination in raw sequencing files is not a good predictor for the amount of human contamination that is retained in the target BAM files after alignment to the target reference genome.

We then estimated mRL and PMD^R^ for the true ancient sequences and the contaminant sequences. For both mammoth and dog samples we found a clear distinction in PMD^R^ of the sequences mapping to the target species and the ones mapped to human, with higher PMD^R^ for the target species, representing true ancient sequences, and lower for the human sequences (Fig. 4C, 4F). However, we found that the contaminant human reads also displayed a lower mRL (Fig. 4B, 4E). This was contrary to the expectation of modern human contaminant sequences being longer than true ancient sequences, but can be explained by the fact that shorter contaminant sequences align easier to evolutionary conserved regions of the target species reference genome than longer sequences (de Filippo, Meyer, and Prüfer 2018; Lee and Schatz 2012).

Considering species by species, the mammoth samples displayed a clearer distinction in PMD^R^ than the dog samples when comparing the reads mapped to the target and to the human parts of the concatenated reference (Fig. 4A, 4D). This may be related to both the age of the samples post-mortem and the age since collection. PMD scores are roughly proportional to the sample’s age (Skoglund et al. 2014), and while the mammoth samples are thousands of years old, they have been housed in collections for less than 30 years. The dog samples on the other hand were only a maximum of 1,000 years old (Supplementary Table 1), but were housed in museum collections since their excavation or collection for up to 125 years. Because the conditions in museum collections are usually far from ideal for DNA preservation (Burrell, Disotell, and Bergey 2015; Díez-del-Molino et al. 2018), this extended period of storage therefore could have had an impact on the preservation of both endogenous dog and contaminant human DNA sequences in the ancient dog samples comparison to the mammoth samples.

### Excluding contaminant reads from faunal BAM files

The presence of contaminant human sequences in ancient faunal BAM files can be challenging for any downstream analyses that are based on evolutionary conserved parts of the genome, such as coding regions, since the contaminant sequences are concentrated in these regions. Other downstream analyses based on genome-wide scans such as estimations of heterozygosity, estimation of inbreeding levels using runs-of-homozygosity, or analyses focused on the presence of rare variants (Schiffels et al. 2016) can be highly affected by the emergence of false variants caused by human contamination (Renaud, Hanghøj, et al. 2019; Llamas et al. 2017). This is especially true for analyses based on low to medium coverage samples, such as most ancient DNA studies. Additionally, since an unknown fraction of the reads discarded using competitive mapping can be of human origin, our detected levels of exogenous human sequences in ancient faunal alignments represent only the lower bound of contamination for these files.

We therefore propose that the method applied here, using competitive mapping of the raw data to a concatenated reference genome composed by the reference genome of the target species and the human genome, represents a fast and simple approach to effectively exclude contaminating human DNA from ancient faunal BAM files (Fig. 2B). An additional advantage of this approach is that a large part of contamination from short microbial reads, common in ancient datasets (de Filippo, Meyer, and Prüfer 2018), should also be excluded with this method as many of these short reads would align to both target and human parts of the concatenated reference and are filtered out using the mapping quality filters.

One relevant downside of using competitive mapping could be the loss of data. True ancient sequences from the target species that belong to conserved regions of the genome and are identical between the target species and human, would align to both parts of the concatenated reference, and thus be lost when using the mapping quality filters. However, our results indicate that the amount of data lost this way is very limited in a genome-wide context (average 1.3%), and slightly concentrated in conserved regions of the genome (average 4.5%). Unfortunately, we do not have a practical way to estimate what fraction of those sequences are true target sequences and how many are of human or microbial origin.

## Conclusions

We show that variable levels of contaminant human sequences exist in ancient faunal datasets. To some extent, this human contamination persists even after sequence reads have been mapped to faunal reference genomes, and is then characterized by short fragment lengths that are concentrated in evolutionary conserved regions of the genome. This results in human contaminant sequences being included in ancient faunal alignment files and thus have the potential to affect a range of downstream analyses. To address this, we here propose a fast and simple strategy: competitive mapping of raw sequencing data to a concatenated reference composed of the target species genome and a human genome, where only the sequences aligned to the target part of the concatenated reference genome are kept for downstream analyses. This approach leads to a small loss of data, but allows for the effective removal of the putative human contaminant sequences.

Contamination is a key issue in ancient DNA studies. Preventive measures both during field collection and in the laboratory therefore remain a critical aspect of ancient DNA research (Llamas et al. 2017; Korlević et al. 2015). There is a growing array of computational methods that allow to confidently identify contamination levels (reviewed in Renaud, Schubert, et al. 2019), but few that allow to efficiently separate authentic ancient sequences from contaminating DNA (Skoglund et al. 2014; de Filippo, Meyer, and Prüfer 2018). Thus, the method we propose here represents an important addition to the selection of tools aimed at computationally reducing the effects of human contamination in ancient faunal DNA research.

## Materials and Methods

### Samples

We analyzed genomic data from 70 ancient and historical mammalian specimens, 50 dogs and 20 woolly mammoths (Supplementary Table 1). The materials derived from dogs originate from a variety of contexts (ethnographic collections and archaeological excavations) and materials (teeth and bones) which have been stored in museum collections for up to 125 years after collection/excavation. The twenty mammoth samples were all collected in Wrangel Island in several expeditions along the last 30 years.

### Laboratory procedures

For all samples, the outer layers of bones, teeth and tusk were removed using an electric powered drill (Dremel, USA) in order to minimize external contamination. Approximately 50 mg of bone powder was recovered from inside the bone, tooth or tusk using an electric drill operated at low speed. We then extracted DNA from all samples using the silica-based protocol described in (Ersmark et al. 2015). Thirty-four of the dog samples were additionally subjected to a pre-digestion step, incubated with EDTA, urea, and proteinase K for one hour at 55°C, to further reduce the amount of contamination within the extract by removing the superficial DNA. We did not treat any of the extracts with USER enzyme in order to enable assessment of post-mortem damage rates following DNA sequencing.

We constructed Illumina genomic libraries for sequencing from the DNA extracts using established ancient DNA protocols (M. Meyer and Kircher 2010; Carøe et al. 2017). All libraries were amplified using indexes unique for each sample and were subsequently pooled and sequenced on a total of 4 lanes on the Illumina HiSeq2500 platform at the National Genomics Infrastructure (Science for Life Laboratory, Stockholm), using paired-end 2×150bp settings.

### Data analyses

We trimmed sequencing adapters and merged paired-end reads using *SeqPrep v*.*1*.*1* (github.com/jstjohn/SeqPrep) with default settings (excluding reads shorter than 30bp) and a slight modification of the source code to calculate the base qualities in the overlapping region (Palkopoulou et al. 2015). We then mapped the merged reads to three separate reference genomes: the African savannah elephant genome (LoxAfr4, Broad Institute), the dog genome (CanFam3.1, Lindblad-Toh et al. 2005), and the human reference genome (Hg19). All mappings were performed using *BWA aln v0*.*7*.*8* (Li and Durbin 2009) using settings adapted for ancient DNA as in Pečnerová et al. (2017).

We removed PCR duplicates from the alignments using a custom script which takes into account both starting and end coordinates of the reads to be identified as duplicates (Palkopoulou et al. 2018) and estimated the number of unique mapping reads using *samtools v1*.*8* (Li et al. 2009). In all cases, we refer to *mapped reads* to those sequences retained after filtering by mapping quality > 30. We consider true endogenous sequences those mapping to the target species (i.e dog reference for ancient dog samples and elephant reference for mammoth samples) and exogenous contaminant sequences those mapping to the non-target reference (i.e elephant and human references for ancient dog samples and dog and human references for mammoth samples). To characterize the sequences mapping to the target reference genome as well as the ones mapping to the non-target and human references using two characteristics of ancient DNA: short fragment size (Rogaev et al. 2006; Allentoft et al. 2012; Kistler et al. 2017) measured as median read length (mRL) and deamination patterns (Briggs et al. 2010; Hofreiter, Jaenicke, et al. 2001) measured as post-mortem damage scores (PMD, (Skoglund et al. 2014). For each sample, we define the PMD ratio (PMD^R^) as the fraction of sequences that display a PMD score > 5. Therefore, a higher PMD^R^ value indicates that the sample contains more sequences with larger PMD scores, thus it is more ‘ancient’.

In order to estimate the amount of data lost using competitive mapping we identified conserved regions between the elephant and human genomes as well as the dog and human genomes. We first used a custom script to split the human reference genome into non-overlapping 35bp long sequences. We then mapped the obtained short sequences to the other two reference genomes, dog and elephant, using *BWA aln* with settings optimized for mapping short reads (Li 2013; Schubert et al. 2012). For each mapping, we filtered out reads with mapping quality below 30 and identified all genomic regions with at least one read mapped. The resulting BED files were used together with *samtools flagstat* to estimate the number of reads mapping to conserved regions before and after competitive mapping.

## Acknowledgements

The authors would like to acknowledge support from Science for Life Laboratory, the National Genomics Infrastructure, and UPPMAX (project numbers: b2014312 and SNIC2020/5-3) for providing assistance in massive parallel sequencing and computational infrastructure. Genetic analyses were funded through a grant from the Swedish Research Council (VR grant 2012-3869) awarded to L.D.. J.v.S. and LD. acknowledge support from FORMAS (project 2015-676), T.R.F. acknowledges support from the EU-funded ITN project ArchSci2020 (grant no. 676154) and for the Qimmeq Project funding from the Velux Foundations, the Aage og Johanne Louis-Hansens Fond and the Wellcome Trust (grant no. 210119/Z/18/Z). D.D.dM. was supported through a Carl Tryggers scholarship (grant CTS17:109).

## Author contributions

T.R.F. and D.D.dM. conceived the study with input from the rest of the coauthors. T.R.F., E.P., and J.vS. performed lab procedures. T.R.F. and D.D.dM. analyzed the data. T.R.F. and D.D.dM. wrote the manuscript with contributions from all other coauthors. All authors contributed to and approved the final version of the manuscript.

## Data availability

All sequencing data generated in this study are available at the European Nucleotide Archive (ebi.ac.uk/ena) with accession numbers XXX-XXX.

## Supplementary Information

**Figure S1:**
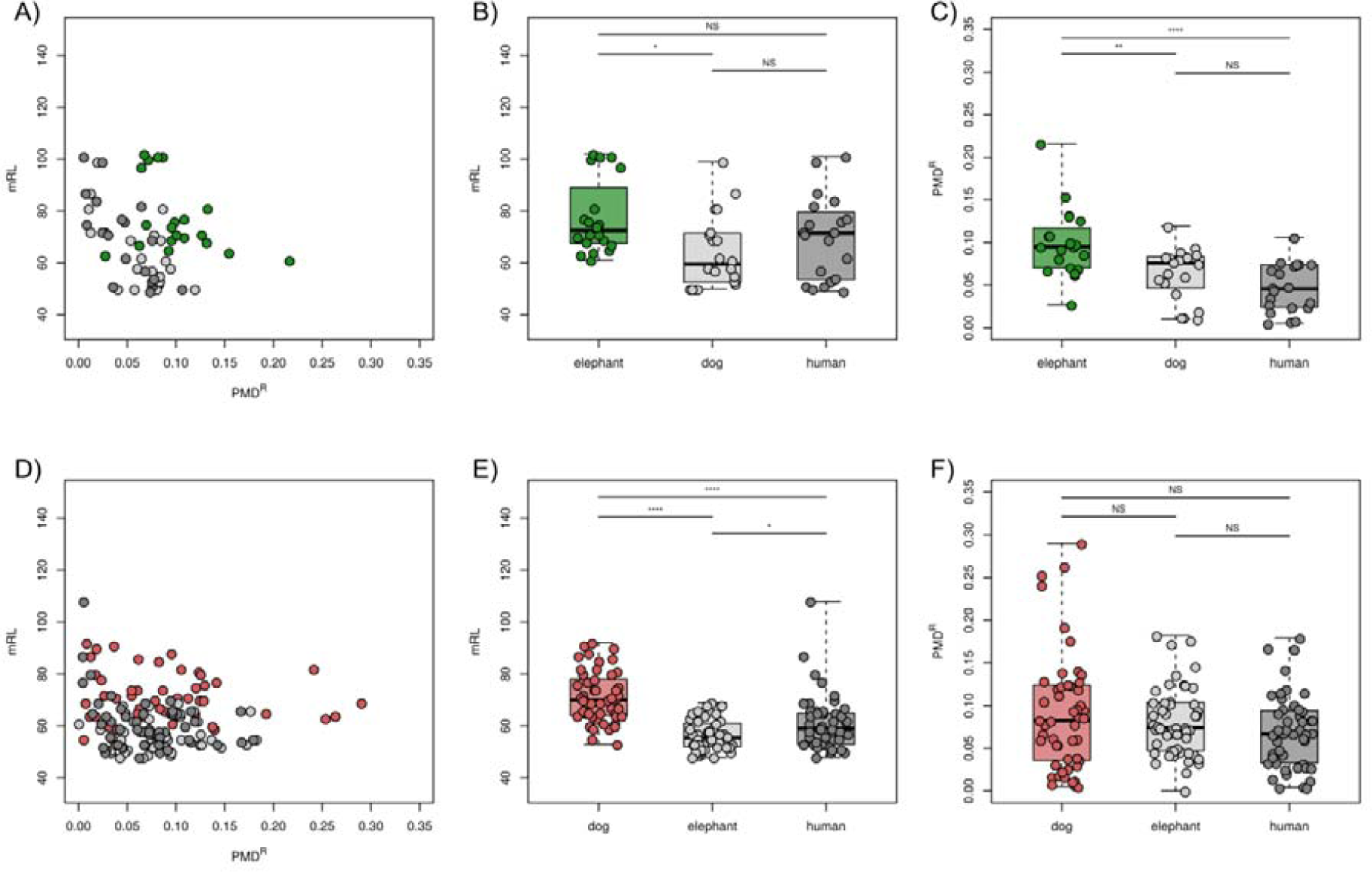
Characterization of sequences mapping to the target, non-target and human references. A) Comparisons of PMD^R^ and mRL for all mammoth samples. B) mRL for mammoth sequences mapping to the elephant, dog and human references. B) PMD^R^ for mammoth sequences mapping to the elephant, dog and human references. D) Comparisons of PMD^R^ and mRL for all ancient dog samples. B) mRL for dog sequences mapping to the elephant, dog and human references. B) PMD^R^ for dog sequences mapping to the elephant, dog and human references. All pairwise comparisons are done using Tukey’s tests. In all cases, NS: p-value >0.05, *: p-value <0.05, **: p-value < 0.01 and ****: p-value < 0.0001.

## References

Allentoft, Morten E., Matthew Collins, David Harker, James Haile, Charlotte L. Oskam, Marie L. Hale, Paula F. Campos, et al. 2012. “The Half-Life of DNA in Bone: Measuring Decay Kinetics in 158 Dated Fossils.” Proceedings of the Royal Society B: Biological Sciences. https://doi.org/10.1098/rspb.2012.1745.

Allentoft, Morten E., Martin Sikora, Karl-Göran Sjögren, Simon Rasmussen, Morten Rasmussen, Jesper Stenderup, Peter B. Damgaard, et al. 2015. “Population Genomics of Bronze Age Eurasia.” Nature 522 (7555): 167–72.

Barlow, Axel, Johanna L. A. Paijmans, Federica Alberti, Boris Gasparyan, Guy Bar-Oz, Ron Pinhasi, Irina Foronova, et al. 2020. “Middle Pleistocene Cave Bear Genome Calibrates the Evolutionary History of Palaearctic Bears.” https://doi.org/10.2139/ssrn.3523359.

Briggs, Adrian W., Udo Stenzel, Philip L. F. Johnson, Richard E. Green, Janet Kelso, Kay Prüfer, Matthias Meyer, et al. 2007. “Patterns of Damage in Genomic DNA Sequences from a Neandertal.” Proceedings of the National Academy of Sciences of the United States of America 104 (37): 14616–21.

Briggs, Adrian W., Udo Stenzel, Matthias Meyer, Johannes Krause, Martin Kircher, and Svante Pääbo. 2010. “Removal of Deaminated Cytosines and Detection of in Vivo Methylation in Ancient DNA.” Nucleic Acids Research 38 (6): e87.

Burrell, Andrew S., Todd R. Disotell, and Christina M. Bergey. 2015. “The Use of Museum Specimens with High-Throughput DNA Sequencers.” Journal of Human Evolution 79 (February): 35–44.

Carøe, Christian, Shyam Gopalakrishnan, Lasse Vinner, Sarah S. T. Mak, Mikkel-Holger S. Sinding, José A. Samaniego, Nathan Wales, Thomas Sicheritz-Pontén, and M. Thomas P. Gilbert. 2017. “Single-Tube Library Preparation for Degraded DNA.” Methods in Ecology and Evolution.

Cooper, A., and H. N. Poinar. 2000. “Ancient DNA: Do It Right or Not at All.” Science. American Association for the Advancement of Science.

Dabney, Jesse, Michael Knapp, Isabelle Glocke, Marie-Theres Gansauge, Antje Weihmann, Birgit Nickel, Cristina Valdiosera, et al. 2013. “Complete Mitochondrial Genome Sequence of a Middle Pleistocene Cave Bear Reconstructed from Ultrashort DNA Fragments.” Proceedings of the National Academy of Sciences of the United States of America 110 (39): 15758–63.

Der Sarkissian, C., L. Ermini, H. Jónsson, A. N. Alekseev, E. Crubezy, B. Shapiro, and L. Orlando. 2014. “Shotgun Microbial Profiling of Fossil Remains.” Molecular Ecology 23 (7): 1780–98.

Der Sarkissian, Clio, Morten E. Allentoft, María C. Ávila-Arcos, Ross Barnett, Paula F. Campos, Enrico Cappellini, Luca Ermini, et al. 2015. “Ancient Genomics.” Philosophical Transactions of the Royal Society of London. Series B, Biological Sciences 370 (1660): 20130387.

Díez-del-Molino, David, Fatima Sánchez-Barreiro, Ian Barnes, M. Thomas P. Gilbert, and Love Dalén. 2018. “Quantifying Temporal Genomic Erosion in Endangered Species.” Trends in Ecology & Evolution 33 (3): 176–85.

Ersmark, Erik, Ludovic Orlando, Edson Sandoval-Castellanos, Ian Barnes, Ross Barnett, Anthony Stuart, Adrian Lister, and Love Dalén. 2015. “Population Demography and Genetic Diversity in the Pleistocene Cave Lion.” Open Quaternary 1 (4): 1–14.

Filippo, Cesare de, Matthias Meyer, and Kay Prüfer. 2018. “Quantifying and Reducing Spurious Alignments for the Analysis of Ultra-Short Ancient DNA Sequences.” BMC Biology 16 (1): 121.

Fu, Qiaomei, Heng Li, Priya Moorjani, Flora Jay, Sergey M. Slepchenko, Aleksei A. Bondarev, Philip L. F. Johnson, et al. 2014. “Genome Sequence of a 45,000-Year-Old Modern Human from Western Siberia.” Nature 514 (7523): 445–49.

Gamba, Cristina, Kristian Hanghøj, Charleen Gaunitz, Ahmed H. Alfarhan, Saleh A. Alquraishi, Khaled A. S. Al-Rasheid, Daniel G. Bradley, and Ludovic Orlando. 2016. “Comparing the Performance of Three Ancient DNA Extraction Methods for High-Throughput Sequencing.” Molecular Ecology Resources 16 (2): 459–69.

Gilbert, M. Thomas P., Eske Willerslev, Anders J. Hansen, Ian Barnes, Lars Rudbeck, Niels Lynnerup, and Alan Cooper. 2003. “Distribution Patterns of Postmortem Damage in Human Mitochondrial DNA.” American Journal of Human Genetics 72 (1): 32–47.

Green, Richard E., Johannes Krause, Adrian W. Briggs, Tomislav Maricic, Udo Stenzel, Martin Kircher, Nick Patterson, et al. 2010. “A Draft Sequence of the Neandertal Genome.” Science 328 (5979): 710–22.

Green, Richard E., Johannes Krause, Susan E. Ptak, Adrian W. Briggs, Michael T. Ronan, Jan F. Simons, Lei Du, et al. 2006. “Analysis of One Million Base Pairs of Neanderthal DNA.” Nature 444 (7117): 330–36.

Hofreiter, M., V. Jaenicke, D. Serre, A. von Haeseler, and S. Pääbo. 2001. “DNA Sequences from Multiple Amplifications Reveal Artifacts Induced by Cytosine Deamination in Ancient DNA.” Nucleic Acids Research 29 (23): 4793–99.

Hofreiter, M., D. Serre, H. N. Poinar, M. Kuch, and S. Pääbo. 2001. “Ancient DNA.” Nature Reviews. Genetics 2 (5): 353–59.

Kircher, Martin. 2012. “Analysis of High-Throughput Ancient DNA Sequencing Data.” Methods in Molecular Biology 840: 197–228.

Kistler, Logan, Roselyn Ware, Oliver Smith, Matthew Collins, and Robin G. Allaby. 2017. “A New Model for Ancient DNA Decay Based on Paleogenomic Meta-Analysis.” Nucleic Acids Research 45 (11): 6310–20.

Korlevic, Petra, Tobias Gerber, Marie-Theres Gansauge, Mateja Hajdinjak, Sarah Nagel, Ayinuer Aximu-Petri, and Matthias Meyer. 2015. “Reducing Microbial and Human Contamination in DNA Extractions from Ancient Bones and Teeth.” BioTechniques 59 (2): 87–93.

Lazaridis, Iosif, Nick Patterson, Alissa Mittnik, Gabriel Renaud, Swapan Mallick, Karola Kirsanow, Peter H. Sudmant, et al. 2014. “Ancient Human Genomes Suggest Three Ancestral Populations for Present-Day Europeans.” Nature 513 (7518): 409–13.

Lee, Hayan, and Michael C. Schatz. 2012. “Genomic Dark Matter: The Reliability of Short Read Mapping Illustrated by the Genome Mappability Score.” Bioinformatics 28 (16): 2097–2105.

Li, Heng. 2013. “Aligning Sequence Reads, Clone Sequences and Assembly Contigs with BWA-MEM.” arXiv [q-bio.GN]. arXiv. http://arxiv.org/abs/1303.3997.

Li, Heng, and Richard Durbin. 2009. “Fast and Accurate Short Read Alignment with Burrows-Wheeler Transform.” Bioinformatics 25 (14): 1754–60.

Li, Heng, Bob Handsaker, Alec Wysoker, Tim Fennell, Jue Ruan, Nils Homer, Gabor Marth, Goncalo Abecasis, Richard Durbin, and 1000 Genome Project Data Processing Subgroup. 2009. “The Sequence Alignment/Map Format and SAMtools.” Bioinformatics 25 (16): 2078–79.

Lindahl, T. 1993. “Instability and Decay of the Primary Structure of DNA.” Nature 362 (6422): 709–15.

Lindblad-Toh, Kerstin, Claire M. Wade, Tarjei S. Mikkelsen, Elinor K. Karlsson, David B. Jaffe, Michael Kamal, Michele Clamp, et al. 2005. “Genome Sequence, Comparative Analysis and Haplotype Structure of the Domestic Dog.” Nature 438 (7069): 803–19.

Llamas, Bastien, Guido Valverde, Lars Fehren-Schmitz, Laura S. Weyrich, Alan Cooper, and Wolfgang Haak. 2017. “From the Field to the Laboratory: Controlling DNA Contamination in Human Ancient DNA Research in the High-Throughput Sequencing Era.” STAR: Science & Technology of Archaeological Research 3 (1): 1–14.

Malmström, Helena, Jan Storå, Love Dalén, Gunilla Holmlund, and Anders Götherström. 2005. “Extensive Human DNA Contamination in Extracts from Ancient Dog Bones and Teeth.” Molecular Biology and Evolution 22 (10): 2040–47.

Meyer, Matthias, Juan-Luis Arsuaga, Cesare de Filippo, Sarah Nagel, Ayinuer Aximu-Petri, Birgit Nickel, Ignacio Martínez, et al. 2016. “Nuclear DNA Sequences from the Middle Pleistocene Sima de Los Huesos Hominins.” Nature, 1–15.

Meyer, Matthias, Martin Kircher, Marie-Theres Gansauge, Heng Li, Fernando Racimo, Swapan Mallick, Joshua G. Schraiber, et al. 2012. “A High-Coverage Genome Sequence from an Archaic Denisovan Individual.” Science 338 (6104): 222–26.

Meyer, M., and M. Kircher. 2010. “Illumina Sequencing Library Preparation for Highly Multiplexed Target Capture and Sequencing.” Cold Spring Harbor Protocols. https://doi.org/10.1101/pdb.prot5448.

Orlando, Ludovic, Aurélien Ginolhac, Guojie Zhang, Duane Froese, Anders Albrechtsen, Mathias Stiller, Mikkel Schubert, et al. 2013. “Recalibrating Equus Evolution Using the Genome Sequence of an Early Middle Pleistocene Horse.” Nature 499: 74–78.

Palkopoulou, Eleftheria, Mark Lipson, Swapan Mallick, Svend Nielsen, Nadin Rohland, Sina Baleka, Emil Karpinski, et al. 2018. “A Comprehensive Genomic History of Extinct and Living Elephants.” Proceedings of the National Academy of Sciences of the United States of America 115 (11): E2566–74.

Palkopoulou, Eleftheria, Swapan Mallick, Pontus Skoglund, Jacob Enk, Nadin Rohland, Heng Li, Ayça Omrak, et al. 2015. “Complete Genomes Reveal Signatures of Demographic and Genetic Declines in the Woolly Mammoth.” Current Biology: CB 25 (10): 1395–1400.

Pecnerová, Patrícia, David Díez-Del-Molino, Nicolas Dussex, Tatiana Feuerborn, Johanna von Seth, Johannes van der Plicht, Pavel Nikolskiy, Alexei Tikhonov, Sergey Vartanyan, and Love Dalén. 2017. “Genome-Based Sexing Provides Clues about Behavior and Social Structure in the Woolly Mammoth.” Current Biology: CB 27 (22): 3505–10.e3.

Prüfer, Kay, Udo Stenzel, Michael Hofreiter, Svante Pääbo, Janet Kelso, and Richard E. Green. 2010. “Computational Challenges in the Analysis of Ancient DNA.” Genome Biology 11 (5): R47.

Racimo, Fernando, Gabriel Renaud, and Montgomery Slatkin. 2016. “Joint Estimation of Contamination, Error and Demography for Nuclear DNA from Ancient Humans.” PLOS Genetics. https://doi.org/10.1371/journal.pgen.1005972.

Rasmussen, Morten, Martin Sikora, Anders Albrechtsen, Thorfinn Sand Korneliussen, J. Víctor Moreno-Mayar, G. David Poznik, Christoph P. E. Zollikofer, et al. 2015. “The Ancestry and Affiliations of Kennewick Man.” Nature 523 (7561): 455–58.

Renaud, Gabriel, Kristian Hanghøj, Thorfinn Sand Korneliussen, Eske Willerslev, and Ludovic Orlando. 2019. “Joint Estimates of Heterozygosity and Runs of Homozygosity for Modern and Ancient Samples.” Genetics 212 (3): 587–614.

Renaud, Gabriel, Mikkel Schubert, Susanna Sawyer, and Ludovic Orlando. 2019. “Authentication and Assessment of Contamination in Ancient DNA.” Methods in Molecular Biology 1963: 163–94.

Rogaev, Evgeny I., Yuri K. Moliaka, Boris A. Malyarchuk, Fyodor A. Kondrashov, Miroslava V. Derenko, Ilya Chumakov, and Anastasia P. Grigorenko. 2006. “Complete Mitochondrial Genome and Phylogeny of Pleistocene mammothMammuthus Primigenius.” PLoS Biology 4 (3). https://journals.plos.org/plosbiology/article/file?type=printable&id=10.1371/journal.pbio.0040073.

Sawyer, Susanna, Johannes Krause, Katerina Guschanski, Vincent Savolainen, and Svante Pääbo. 2012. “Temporal Patterns of Nucleotide Misincorporations and DNA Fragmentation in Ancient DNA.” PloS One 7 (3): e34131.

Schiffels, Stephan, Wolfgang Haak, Pirita Paajanen, Bastien Llamas, Elizabeth Popescu, Louise Loe, Rachel Clarke, et al. 2016. “Iron Age and Anglo-Saxon Genomes from East England Reveal British Migration History.” Nature Communications 7 (January): 10408.

Schubert, Mikkel, Aurelien Ginolhac, Stinus Lindgreen, John F. Thompson, Khaled A. S. Al-Rasheid, Eske Willerslev, Anders Krogh, and Ludovic Orlando. 2012. “Improving Ancient DNA Read Mapping against Modern Reference Genomes.” BMC Genomics 13 (May): 178.

Skoglund, Pontus, Bernd H. Northoff, Michael V. Shunkov, Anatoli P. Derevianko, Svante Pääbo, Johannes Krause, and Mattias Jakobsson. 2014. “Separating Endogenous Ancient DNA from Modern Day Contamination in a Siberian Neandertal.” Proceedings of the National Academy of Sciences of the United States of America 111 (6): 2229–34.

Slatkin, Montgomery, and Fernando Racimo. 2016. “Ancient DNA and Human History.” Proceedings of the National Academy of Sciences 2016 (23): 1–8.

Smith, T. F., M. S. Waterman, and C. Burks. 1985. “The Statistical Distribution of Nucleic Acid Similarities.” Nucleic Acids Research 13 (2): 645–56.

Stiller, M., R. E. Green, M. Ronan, J. F. Simons, L. Du, W. He, M. Egholm, et al. 2006. “Patterns of Nucleotide Misincorporations during Enzymatic Amplification and Direct Large-Scale Sequencing of Ancient DNA.” Proceedings of the National Academy of Sciences of the United States of America 103 (37): 13578–84.

Willerslev, Eske, and Alan Cooper. 2005. “Ancient DNA.” Proceedings. Biological Sciences / The Royal Society 272 (1558): 3–16.

